# No evidence that plasmablasts transdifferentiate into developing neutrophils in severe COVID-19 disease

**DOI:** 10.1101/2020.09.27.312538

**Authors:** José Alquicira-Hernandez, Joseph E Powell, Tri Giang Phan

**Affiliations:** Garvan Institute of Medical Research, 384 Victoria St, Darlinghurst NSW 2010, Sydney, Australia; UNSW Cellular Genomics Futures Institute, University of New South Wales, Sydney, Australia; St Vincent’s Clinical School, Faculty of Medicine, UNSW Sydney, 384 Victoria St, Darlinghurst NSW 2010, Sydney, Australia

## Abstract

A recent study by Wilk et al. of the transcriptome of peripheral blood mononuclear cells (PBMCs) in seven patients hospitalized with COVID-19 described a population of “developing neutrophils” that were “phenotypically related by dimensionality reduction” to plasmablasts, and that these two cell populations represent a “linear continuum of cellular phenotype”^1^. The authors suggest that, in the setting of acute respiratory distress syndrome (ARDS) secondary to severe COVID-19, a “differentiation bridge from plasmablasts to developing neutrophils” connected these distantly related cell types. This conclusion is controversial as it appears to violate several basic principles in cell biology relating to cell lineage identity and fidelity. Correctly classifying cells and their developmental history is an important issue in cell biology and we suggest that this conclusion is not supported by the data as we show here that: (1) regressing out covariates such as unique molecular identifiers (UMIs) can lead to overfitting; and (2) that UMAP embeddings may reflect the expression of similar genes but not necessarily direct cell lineage relationships.

## Main

Infection-induced plasmablasts are proliferating plasma cells derived from B cells of the lymphoid cell lineage that secrete antibodies against invading pathogens^2^. In contrast, neutrophils are cells derived from the myeloid cell lineage that trap, phagocytose and kill pathogenic organisms by generating reactive oxygen species, secreting proteases and degradative enzymes and casting neutrophil extracellular traps (NETs)^3^. Accordingly, the lineage conversion of plasmablasts into developing neutrophils requires a massive re-organization of the cell at the level of the transcriptome, epigenome, and proteome. While the forced transdifferentiation of B cell lines, B cell progenitors and naïve B cells into macrophages have been described, notably via the ectopic expression of CCAAT-enhancer-binding protein (C/EBP) family of transcription factors^4,5^, the spontaneous transdifferentiation of terminally differentiated plasma cells is an entirely different hypothesis that would require compelling evidence, which was not presented in the paper.

The recent introduction of high-throughput single cell RNA-sequencing (scRNA-seq) has created new opportunities for developmental biologists to map the developmental history of differentiated cell types^6^. These technologies assay the expression of genes in a large number of cells – their cell transcriptional state – and a standard computational approach is to flatten this high-dimensional data into a two-dimensional (2D) Euclidean space as state manifolds. In these projections, the proximity of one cell type to another may denote similarities in the cell transcriptional state. A major challenge in the analysis of scRNA-seq data is to remove noise from technical variation while preserving biological heterogeneity^7^. To understand how this trade-off is balanced, we first used their code to reproduce figure 1c from their paper (**Fig. 1a**). In doing so, we noted that, in addition to regressing out mitochondrial genes, ribosomal RNA and ribosomal genes, the authors also regressed out the number of UMIs (nCount_RNA) and the number of expressed genes (nFeature_RNA) from the gene expression data in their analysis. This step is unnecessary in Seurat when using the SCTransform normalization method as the number of UMIs is explicitly modelled using a regularised negative binomial regression^7^. Regressing out the number of UMIs using a standard generalised linear model (GLM) is discouraged by the developers due to overfitting^7^. Moreover, the number of expressed genes in each cell is correlated to its number of UMIs (Pearson correlation = 0.9249). We therefore re-analysed the data in three ways: without regressing out any covariates; regressing out only mitochondrial genes (**Fig. 1b**); and regressing out mitochondrial and ribosomal genes (**Fig. 1c**). This exercise shows that the reported relationship between developing neutrophils and plasmablasts breaks down when technical noise is not removed at the expense of biological variability and the data is not overfitted.

**Figure 1.**
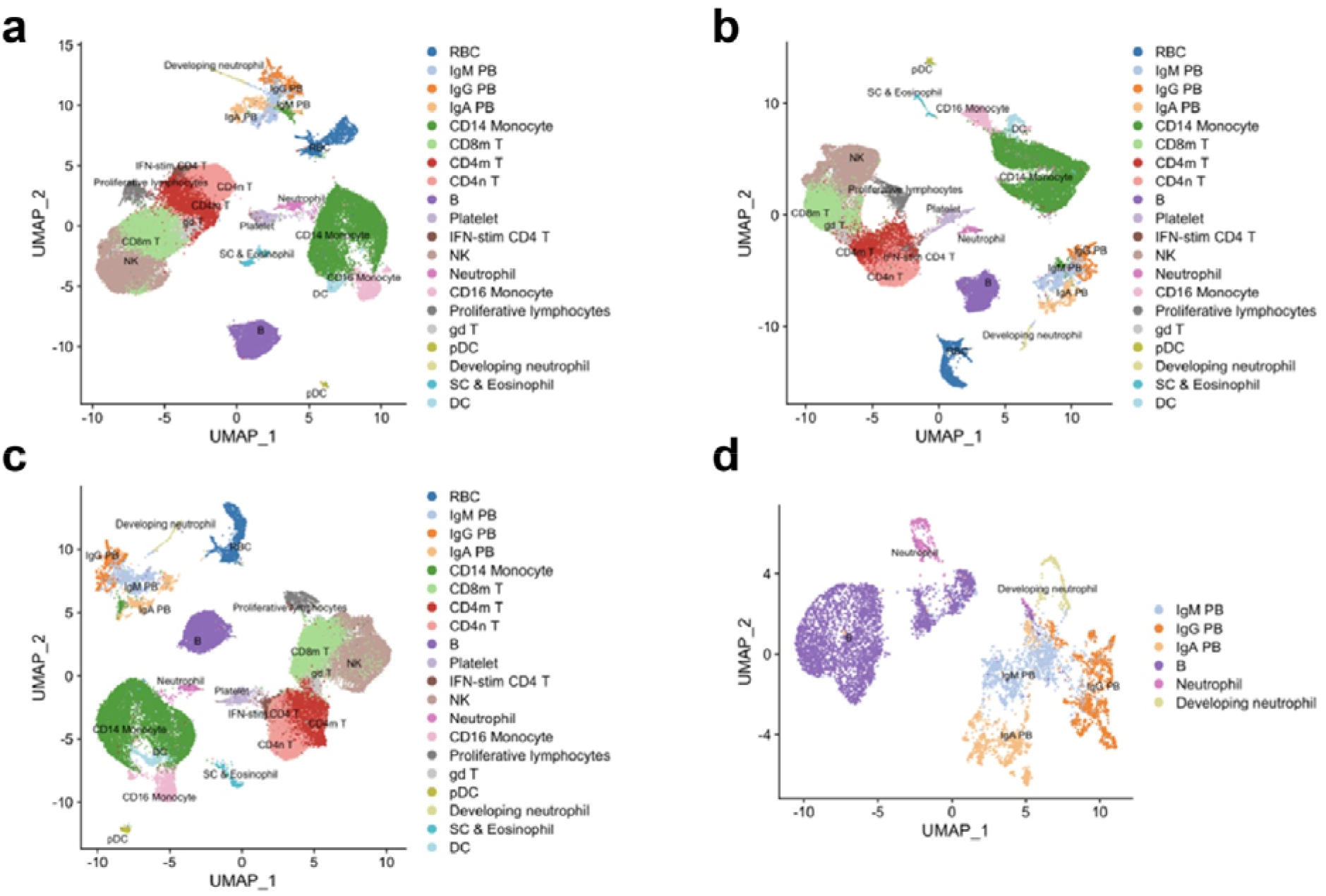
Re-analysis of scRNA-seq data from Wilk et al. show developing neutrophils do not transdifferentiate from plasmablasts. **a.** Reconstruction of the UMAP embedding from figure 1c of Wilk et al. **b.** UMAP embedding showing effect of regressing out mitochondrial genes. **c.** UMAP embedding showing effect of regressing out mitochondrial genes and ribosomal genes. **d.** UMAP embedding of neutrophils, developing neutrophils, B cells and plasma cells.

Accordingly, we believe that orthogonal approaches, such as single cell DNA sequencing to detect rearranged immunoglobulin heavy and light chain variable genes in the developing neutrophils, are needed to provide evidence of their B cell origin and support the conclusion that terminally differentiated plasma cells transdifferentiate into developing neutrophils. Furthermore, unless the data visualization parameters are carefully chosen, the 2D UMAP embeddings may give the misleading impression that cell clusters are closer or further than they actually are. For example, in figure 1c of Wilk et al., developing neutrophils appear to occupy a similar manifold space as plasmablasts. However, from this viewpoint the plasmablasts appear to be distantly related to the B cells from which they are derived. In addition, these types of analyses make the *a priori* assumption that all the cell types being studied are developmentally related and linked by a cell lineage tree. This may hold true when analysing *in vitro* differentiated cells that share a common ancestor cell of origin that is present in the cell culture, but may not apply to PBMCs that contain a heterogeneous collection of cell types that have arisen from different committed progenitors.

Since plasmablasts are derived from B cells, we decided to subset neutrophils, developing neutrophils, B cells and plasmablasts to further explore any similarities between this cell clusters. This analysis revealed that, counter-intuitively, neutrophils were not related to developing neutrophils, and B cells were not related to plasmablasts (**Fig. 1d**). This exercise suggests instead that the developing neutrophils and plasmablasts, in responding to severe COVID-19 disease, are activated cell types that may share the expression of a number of genes and gene modules that have led to their misclassification as related cell types. These gene modules may result from hyperactivation of the immune system by SARS-CoV-2 and reflect the heightened state of cellular activation. Indeed, in their analysis the authors point out that the developing neutrophils were unlikely to be doublets and also unlikely to be granulocytes that had phagocytosed B cells as there were no clinical features of hemophagocytic lymphohistiocytosis (HLH). However, HLH is often difficult to diagnose clinically and severe COVID-19 disease has been linked in some cases with secondary HLH^8,9^.

Nevertheless, the paper by Wilk et al. is a timely and valuable contribution to the understanding of how severe COVID-19 disease reshapes the distribution and activation state of cell clusters in PBMCs. Neutrophils have a high density and are not normally present in Ficoll preparations of blood used to isolate PBMCs. The developing neutrophils may therefore be similar to the pro-inflammatory low-density granulocytes that have been described in a number of autoimmune diseases^10^. These low-density neutrophils consists of hyposegmented immature neutrophils (band forms) that are also produced during emergency granulopoiesis^11^. Thus, it is more plausible that the developing neutrophils are produced in the bone marrow from myeloid precursors in response to overwhelming infection with SARS-CoV-2.

In summary, scRNA-seq is becoming an increasingly popular tool for dissecting cellular heterogeneity in complex biological systems. However, it does have its limitations and, in the example shown here, may mislead and generate interesting hypotheses that risk being erroneously accepted as conclusions without further cross-validation. Our analysis of the data from Wilk et al. indicate that there is not enough evidence to suggest that developing neutrophils transdifferentiate from plasmablasts.

## Author contributions

JAH performed the bioinformatic analysis; JAH, JEP and TGP wrote the manuscript.

## Acknowledgments

JEP is supported by NHMRC Investigator Grant 1175781; TGP is supported by NHMRC Senior Research Fellowship 1155678 and Project Grants 1124681 and 1139865.

